# A cutting-edge approach unravels a novel role for CDK6 in leukemic progenitor cells

**DOI:** 10.1101/2020.10.05.325886

**Authors:** Eszter Doma, Isabella Maria Mayer, Tania Brandstoetter, Barbara Maurer, Reinhard Grausenburger, Ingeborg Menzl, Markus Zojer, Andrea Hoelbl-Kovacic, Leif Carlsson, Gerwin Heller, Karoline Kollmann, Veronika Sexl

## Abstract

Studies of molecular mechanisms of hematopoiesis and leukemogenesis are hampered by the unavailability of progenitor cell lines that accurately mimic the situation *in vivo*. We now report a robust method to generate and maintain LSK (lin^-^, Sca-1^+^, c-Kit^+^) cells which closely resemble MPP1 cells. HPC^LSK^ reconstitute hematopoiesis in lethally irradiated recipient mice over more than eight months. Upon transformation with different oncogenes including BCR/ABL, FLT3-ITD or MLL-AF9 their leukemic counterparts maintain stem cell properties *in vitro* and recapitulate leukemia formation *in vivo*. The method to generate HPC^LSK^ can be applied to transgenic mice and we illustrate it for CDK6-deficient animals. Upon BCR/ABL^p210^ transformation, *Cdk6^-/-^* HPC^LSKs^ induce disease with a significantly enhanced latency and reduced incidence, showing the importance of CDK6 in leukemia formation. Studies of the CDK6 transcriptome in murine HPC^LSK^ and human BCR/ABL^+^ cells have verified that certain pathways depend on CDK6 and have uncovered a novel CDK6-dependent signature, suggesting a role for CDK6 in leukemic progenitor cell homing. Loss of CDK6 may thus lead to a defect in homing. The HPC^LSK^ system represents a unique tool for combined *in vitro* and *in vivo* studies and enables the production of large quantities of genetically modifiable hematopoietic or leukemic stem/progenitor cells.

**Key points:**

1. We describe the generation of murine cell lines (HPC^LSK^) which reliably mimic hematopoietic/leukemic progenitor cells.
2. *Cdk6^-/-^* BCR/ABL^p210^ HPC^LSKs^ uncover a novel role for CDK6 in homing.

## INTRODUCTION

Adult hematopoietic stem cells (HSCs) represent 0.01-0.005% of all nucleated cells in the bone marrow (BM). They are unique in their ability to continuously self-renew, differentiate into distinct lineages of mature blood cells^1^ and regenerate a functional hematopoietic system following transplantation into immunocompromised mice^2–5^. Most hematopoietic malignancies originate in stem/progenitor cells upon acquirement of genetic/epigenetic defects. These so called leukemic stem cells (LSCs) maintain key characteristics of regular HSCs, including the ability of self-renewing and multi-potency^6,7^.

Although hematopoietic cell differentiation is a dynamic and continuous process, cell surface marker expression defining distinct subsets and developmental stages is an inevitable tool in HSC characterization. A common strategy is to further define murine lineage negative, c-Kit and Sca-1 positive (LSK) cells by their CD48, CD135, CD150 and CD34 expression. This marker combination stratifies the most dormant HSCs into the increasingly cycling multipotent progenitors (MPP) 1 and 2 and the myeloid or lymphoid prone MPP3 and 4^8^. Leukemia, analogous to normal hematopoiesis, is hierarchically organized; LSCs residing in the BM initiate and maintain the disease and give rise to their more differentiated malignant progeny. Therapeutically, LSCs are often resistant against many current cancer treatments and thus cause disease relapse^9–13^. Understanding potential Achilles’ heels in LSCs to develop new curative therapeutic approaches is of fundamental interest and represents a major frontier of cancer biology.

Understanding hematopoietic disease development and defining therapeutic intervention sites requires the availability of multi-potential hematopoietic cell lines. HSCs can be maintained and expanded to a very limited extent *in vitro* - the vast majority of their progeny differentiates in culture. Numerous attempts have been made to increase the number of long-term (LT)-HSCs in culture including the use of high levels of cytokines and growth factors or ill-defined factors secreted by feeder cells^14–28^.

Alternatively, immortalization using genetic manipulation was employed to establish stem cell-like cell lines. One major limitation of these cell lines is the failure to reconstitute a fully functional hematopoietic system upon transplantation^29–30^. One of the most successful immortalized murine multipotent hematopoietic cell lines is the EML (Erythroid, Myeloid, and Lymphocytic) line derived by retroviral expression of a truncated, dominant-negative form of the human retinoic acid receptor. However, EML cells are phenotypically and functionally heterogeneous and display a block in the differentiation of myeloid cells^31–38^.

An alternative route for immortalization of murine multipotent hematopoietic cells was employing *Lhx2*^39–41^, a LIM-homeobox domain transcription factor binding a variety of transcriptional co-factors. *Lhx2* is expressed in embryonic hematopoietic locations such as the aorta-gonad-mesonephros (AGM) region, yolk sac and fetal liver, but is absent in BM, spleen and thymus of adult mice^42–44^. *Lhx2* up-regulates key transcriptional regulators for HSCs including *Hox* and *Gata* while down-regulating differentiation-associated genes^39^. *Lhx2* is aberrantly expressed in human chronic myelogenous leukemia suggesting a role for *Lhx2* in the growth of immature hematopoietic cells^45^. Enforced expression of *Lhx2* in BM-derived murine HSCs and embryonic stem cells (ES)/induced pluripotent (iPS) cells resulted in *ex vivo* expansion of engraftable HSC-like cells^41,42,46^ strictly dependent on stem cell factor (SCF) and yet undefined autocrine loops providing additional secreted molecule(s)^40^. These cells generate functional progeny and long term repopulate stem cell–deficient hosts^40,43,47–48^. The cyclin-dependent kinase 6 (CDK6) has been recently described as a critical regulator of HSC quiescence and is essential in BCR/ABL^p210^ LSCs^49–50^. Besides its main characteristic, CDK6 and its close homolog CDK4 control cell cycle progression, CDK6 functions as a transcriptional regulator^51–53^. CDK6 is recognized as being a key oncogenic driver in hematopoietic malignancies and therefore represents a promising target for cancer therapy and intervention^49,54–56^. More recent evidence highlights the importance of CDK6 during stress, including oncogenic transformation when CDK6 counteracts p53 effects^57^. Furthermore, CDK6 plays a crucial role in several myeloid diseases, including Jak2^V617F+^ MPN, CML and AML by regulating stem cell quiescence, apoptosis, differentiation and cytokine secretion^49,56,58–59^.

Using the long term culture system, it was possible to generate HPC^LSKs^ from the transgenic mouse line *Cdk6^-/-^* which represents a powerful tool to analyze specific functions of CDK6 in progenitor cells and allows mechanistic and therapeutic studies tailored specifically to leukemic stem/progenitors cells.

## MATERIALS AND METHODS

### Animals

Mice (C57BL/6N, NSG [NOD.Cg-Prkdcscid Il2rgtm 1Wjl/SzJ], Ly5.1^+^ [B6.SJL-Ptprca]) and *Cdk6^-/-^*^60^ were bred and maintained under special pathogen-free (SPF) conditions at the Institute of Pharmacology and Toxicology, University of Veterinary Medicine, Vienna, Austria. Age-matched (7-11 weeks) male and female mice were used unless indicated otherwise. All procedures were approved by the institutional ethics and animal welfare committee (BMWFW-68.205/0093-WF/V/3b/2015 and BMWFW-68.205/0112-WF/V/3b/2016) and the national authority according to §§26ff. of the Animal Experiment Act, Tierversuchsgesetz 2012 - TVG 2012.

### HPC^LSK^ cell line generation

BM of two to five C57BL/6 mice was isolated, pooled and sorted for LSK cells. Sorted LSK cells were cultured in 48-well-plates for 48 hours in a 1:1 ratio of Stem Pro-34 SFM (Gibco/ Thermo Scientific, Waltham, MA, USA) and Iscoves modified Dulbecco medium (IMDM, Sigma-Aldrich, St. Louis, MO, USA) supplemented with 0.75□×□10^−4^ M 1-Thiolglycerol (MTG, Sigma), Penicillin/Streptomycin (P/S, Sigma), 2 mM L-Glutamine (L-Glut, Sigma), 25 U heparin (Sigma), 10 ng fibroblast growth factor (mFGF) acidic (R&D Sytems, Minneapolis, USA), 10 ng mIGF-II (R&D), 20 ng mTPO (R&D), 10 ng mIL-3 (R&D), 20 ng hIL-6 (R&D) and stem fell factor (SCF, generated in-house) used at 2% final concentration. LSK cells were transduced with a *Lhx2* pMSCV-puromycin (Clontech/Takara, Mountain View, CA, USA) vector^47^ in 1% pegGOLD Universal Agarose (Peqlab/VWR Darmstadt, Germany) coated 48-well-plates and transfected four times on day three to six with the *Lhx2-* containing viral supernatant. At day seven, cells were transferred to 1% agarose-coated 24-well-plates in IMDM with 5% FCS, 1.5□×□10^-4^ M MTG, P/S, 2 mM L-Glut referred hereafter as IMDM culture medium. Additionally, the IMDM culture medium was supplemented with 12.5 ng/ml IL-6 (R&D) and 2% SCF. At day ten, 1.5 μg/ml puromycin (InvivoGen, San Diego, USA) was added to the medium to select for the *Lhx2* expressing LSK cells. The same reagents were subsequently used for all the experiments.

### HPC^LSK^ cell line culture

HPC^LSK^ cell lines were kept on 1% agarose coated culture plates. Solidified plates were stored in a 5% CO2 humidified incubator with 1 ml IMDM culture media per well. HPC^LSK^ cells were plated in IMDM culture media supplemented with 12.5 ng/ml IL-6, 2% SCF and 1.5 μg/ml puromycin on the agarose plates. Cells were continuously kept at a density between 0.8-2□×□10^6^ cells/ml. BM-derived HPC5 cell line was kept in IMDM culture media supplemented with 12.5 ng/ml IL-6 and 2% SCF, while BM-derived HPC9 cells and ES cell line-derived HPC-7 were cultured in IMDM supplemented with 2% SCF (all lines provided by Leif Carlsson). The virus packaging cell lines Platinum-E (Plat-E,Cell Biolabs, Inc, San Diego, CA, USA) and GP^p210^-GFP^61^ were kept in DMEM (Sigma) supplemented with 10% FCS and P/S. The pre-pro-B-BCR/ABL^p185^-GFP were cultured in RPMI (Sigma) with 10% FCS and P/S^57^.

## RESULTS

### Generation of murine hematopoietic progenitor HPC^LSK^ cell lines

To meet the increasing need of studying hematopoietic stem/progenitor cells, we sought to establish a robust method to generate murine stem-cell lines by modifying a strategy that was originally described by the Carlsson lab^41,47^. Sorted murine LSK cells were maintained in cytokine- and growth factor-supplemented serum-free medium for 2 days. Thereafter, the cells were infected with a retroviral construct encoding *Lhx2* coupled to a puromycin selection marker and switched to SCF, IL-6 and 5% serum-containing IMDM medium on agarose-coated plates to prevent attachment-induced differentiation. Puromycin selection was initiated ten days after sorting. Within four weeks continuously proliferating, HPC^LSK^ cell lines establish and can be stored long term by cryopreservation (Fig. 1a). LSK cells can be classified into dormant HSCs and four subsequent MPP populations based on their surface markers^8^. HPC^LSK^ cell lines express c-Kit and Sca-1 but lack expression of the myeloid and lymphoid lineage markers Gr-1 (neutrophil), CD11b (monocyte/macrophage), CD3 (T cell), CD19 (B cell) and Ter119 (erythroid). According to the CD34, CD48 and CD150 expression, HPC^LSKs^ categorize as MPP2 – a population able to give rise to myeloid and lymphoid cells^8^. Despite the MPP2 surface expression markers, transcriptome analysis of HPC^LSKs^ revealed a predominant overlap with the MPP1 signature pointing to an even more immature state. Upon long-term culture a uniform cellular morphology is maintained within the cell lines (Fig 1b, Supplementary Fig. S1a-d). Comparison to other progenitor cell lines including the bone marrow derived BM-HPC5, BM-HPC9 and the ES-derived HPC-7 cell line^41^ showed that HPC^LSKs^ have the most immature profile. The other cell lines are either positive for lineage markers or lack Sca-1 expression. The ES-derived HPC-7 cell line stains positive for c-Kit, Sca-1, CD48 and CD150 and lacks lineage markers. It is also limited in its differentiation capacity^62–63^ (Supplementary Fig. 1e).

**Figure 1:**
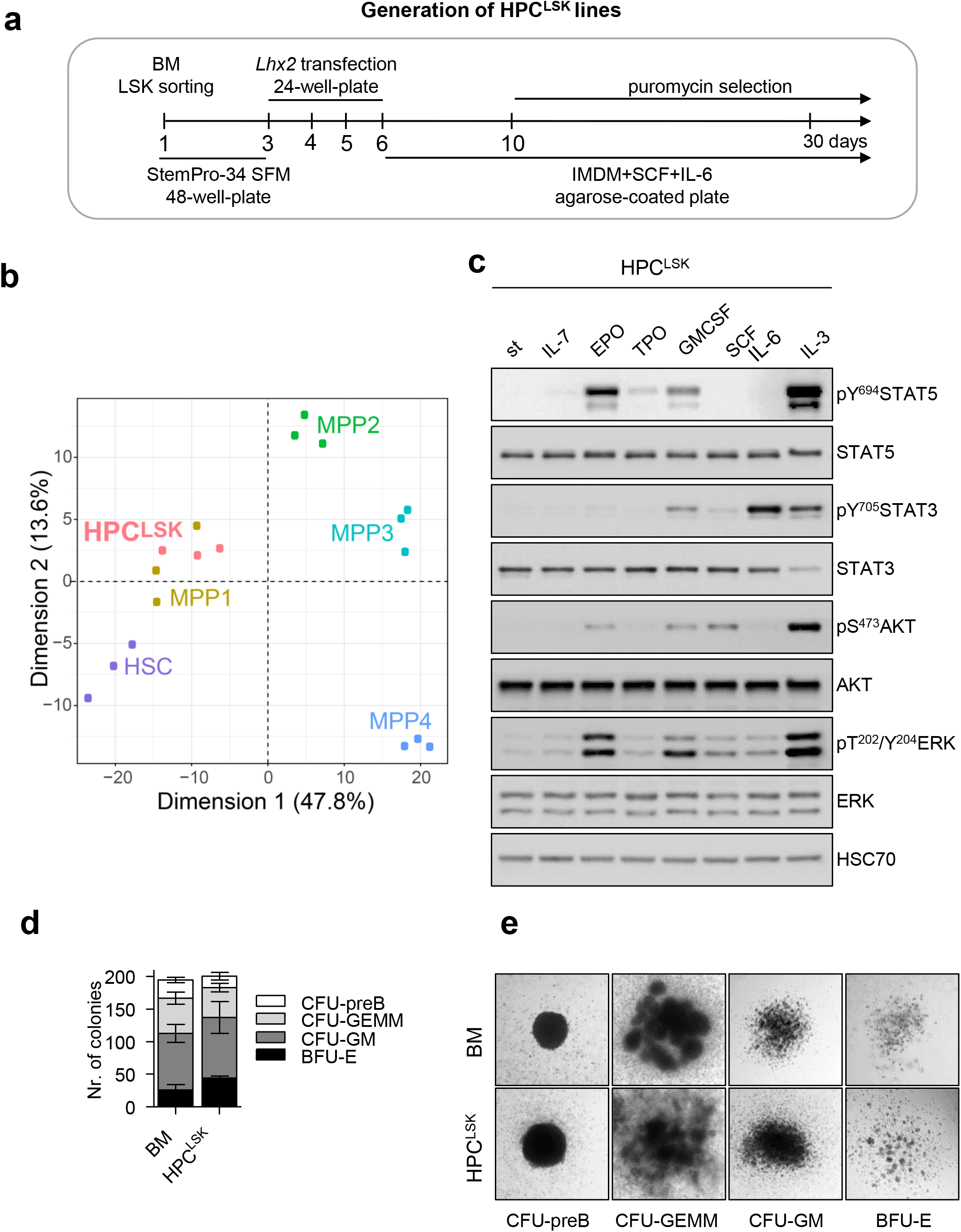
Establishing murine hematopoietic progenitor HPC^LSK^ lines. (**a**) Schematic workflow of HPC^LSK^ cell line establishment. LSKs were sorted from murine BM, transfected with *Lhx2* including a puromycin selection marker and kept in SCF and IL-6 on agarose-coated plates. StemPro-34 SFM: serum free media, IMDM: Iscove’s Modified Dulbecco’s Media, SCF: stem cell factor. (**b**) Principal component analysis of the expression profiles of HPC^LSKs^ (n=3) compared to murine HSCs (batch-corrected top500 variance genes are plotted). (**c**) Immunoblot of lysates from 3h-starved HPC^LSK^ cells followed by treatment with IL-7, EPO, TPO, GMCSF, SCF, IL-6 or IL-3 (100ng/ml each) for 15 min. The presence of total and phosphorylated STAT5, STAT3, AKT and ERK was detected. HSC70 serves as a loading control. st: starved. A representative blot of two independent experiments is shown. (**d**) Colonies with different morphologies were counted. Seeding density of 1 250 HPC^LSKs^ or 240 000 BM cells/35-mm-dish. Error bars represent mean±SD, n≥3. (**e**) Images of colonies formed by HPC^LSK^ cells 10 days after cytokine cocktail treatment (EPO, GMCSF, IL-7, SCF, IL-6, IL-3) in semi-solid methylcellulose gels. BFU-E: burst-forming unit-erythroid, CFU-GM: colony-forming unit-granulocyte macrophage, CFU-GEMM: CFU-granulocyte erythrocyte monocyte megakaryocyte.

### HPC^LSK^ cells are able to differentiate to myeloid and lymphoid cells *in vitro*

To explore signaling patterns, HPC^LSK^ cells were treated with cytokines for 15 min. EPO, GM-CSF or IL-3 resulted in phosphorylation and activation of STAT5, STAT3, AKT and ERK signaling, while IL-6 induced predominantly STAT3 phosphorylation. STAT3, AKT and ERK were also activated upon SCF treatment albeit to a lesser extent in line with signaling in stem/progenitor cells (Fig. 1c). In line, HPC^LSK^ cells formed erythroid (BFU-E), myeloid (CFU-GM, CFU-GEMM) and pre-B (CFU-preB) cell colonies in methylcellulose enriched cytokines (EPO, GMCSF, IL-7, SCF, IL-6, IL-3) comparable to primary BM-derived cells (Fig. 1d-e). We confirmed expression of erythroid (Ter119/CD71), myeloid (CD11b/Gr-1) or B cell (B220/CD93) markers on these colonies (Supplementary Fig 1g). In comparison, the ability to form colonies and to *in vitro* differentiate of HPC-7 and BM-HPC5 cells was reduced in accordance with an impaired cytokine–induced activation of STAT5, STAT3, AKT and ERK (Supplementary Fig. S1h-j).

### HPC^LSKs^ are multipotent *in vivo*

As HPC^LSKs^ differentiate into myeloid and lymphoid lineages *in vitro*, we explored the potential of the cells to protect mice from radiation-induced death *in vivo*. Lethally irradiated Ly5.1^+^ mice received 1×10^7^ Ly5.2^+^ BM-HPC5 or HPC^LSK^ cells per tail vein injection. Ly5.2^+^ BM cells were used as controls. Non-injected irradiated mice died within 10 days, briefly thereafter followed by BM-HPC5 recipients. Injection of HPC^LSKs^ and injection of primary BM cells rescued the mice due to the efficient repopulation of the hematopoietic system (Fig. 2a-b). After 40 days, white blood cell (WBC) and red blood cell (RBC) counts were comparable between HPC^LSKs^–injected and BM-injected controls (Fig. 2c). Blood counts remained stable over a 6-months-period after which the experiment was terminated (Supplementary Fig. S2a). HPC^LSKs^ had efficiently homed to the BM, blood, spleen and thymus comparable to the BM control and no alterations of the spleen weight was detectable (Fig. 2d-e). FACS analysis confirmed the efficient repopulation of the hematopoietic system. Numbers of myeloid and lymphoid progenitors in the BM and differentiated blood cells (Gr-1^+^ granulocytes, CD11b^+^ monocytes, Gr-1/CD11b^+^ eosinophils/neutrophils and B220^+^ B cells) were comparable to BM-injected mice. Only HPC^LSK^-derived CD4^+^ or CD8^+^ T cells were significantly lower in the blood, however, were present in the thymus in similar numbers as in the BM-injected control (Fig. 2f).

**Figure 2:**
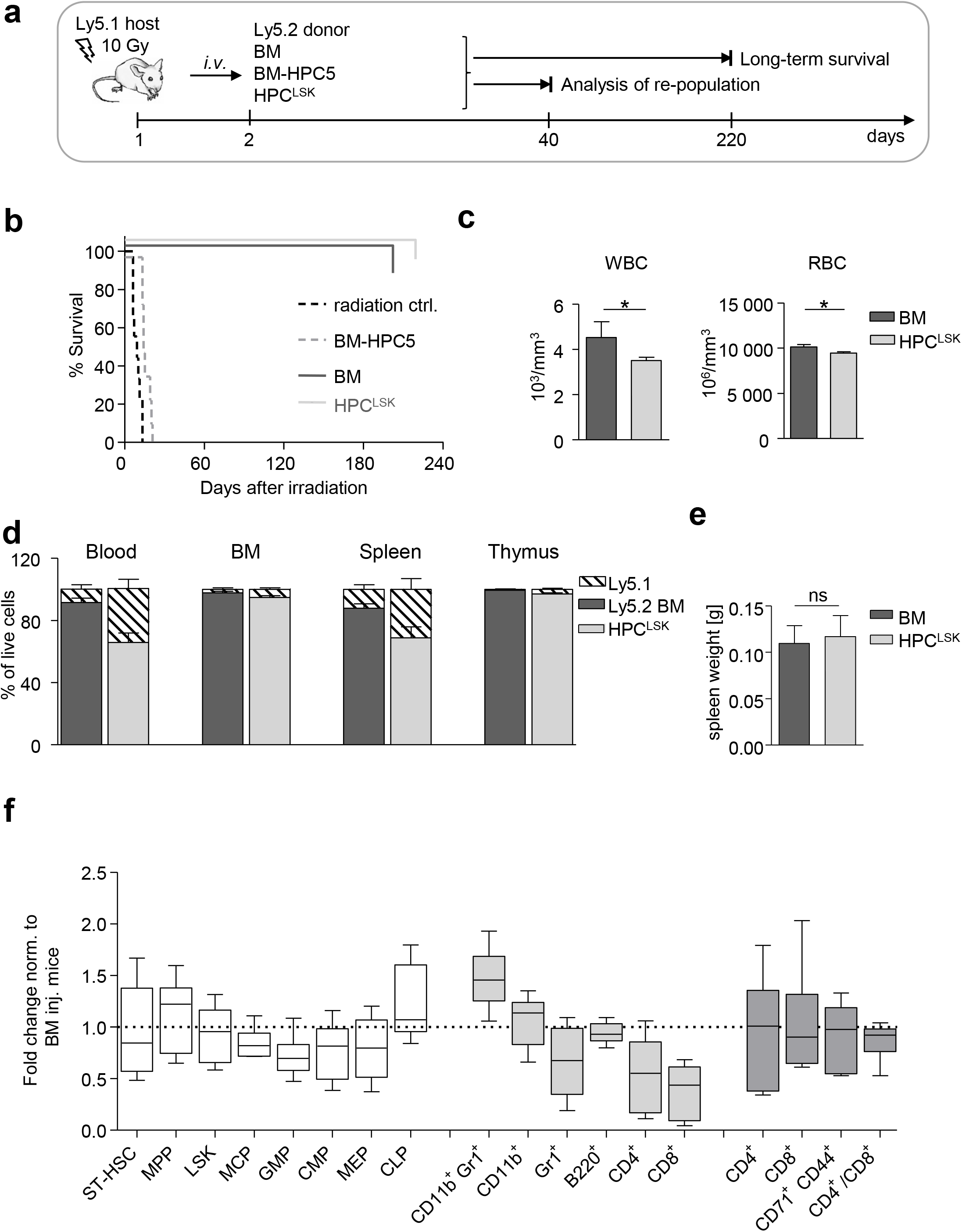
HPC^LSK^ cell lines can repopulate the hematopoietic system. (**a**) Experimental scheme: Ly5.1^+^ recipient mice were lethally irradiated (10 Gy) 24 h prior to *i.v*. injection of 1×10^7^ Ly5.2^+^ BM (positive control), BM-HPC5 or HPC^LSK^ cells. 40 days later, some mice of BM- and HPC^LSK^-injected group were terminated and hematopoietic organs were analyzed. The remaining injected mice were analyzed for their long-term survival. (**b**) Survival of BM- (n=7), BM-HPC5- (n=8) and HPC^LSKs^- (n=10) injected mice compared to irradiation control (n=9), Log-rank (Mantel-Cox) Test \*\*\**P*<0.0001. (**c**) WBC and RBC in peripheral blood of BM- and HPC^LSK^-injected recipients were compared 40 days after treatment. Data are presented as mean±SEM (\**P*<0.01, Student t-test or Mann Whitney test for platelets) in 6-12 mice/group. (**d**) Comparison of Ly5.2^+^ BM versus HPC^LSK^ cells’ engraftment in the blood, BM, spleen and thymus of lethally irradiated Ly5.1^+^ mice after 40 days. Data are presented as mean±SD, n≥4. (**e**) Spleen weights of mice 40 days after lethally irradiation and BM- or HPC^LSK^-injection. Data represent mean±SD, n≥5. (**f**) Composition of the engrafted Ly5.2^+^ HPC^LSK^ cells in blood, BM and thymus after 40 days. ST-HSC; MPP (Lin^-^, Sca-1^+^, c-Kit^+^, CD150^-^, CD48^+^), LSKs, MCP (myeloid committed progenitor, Lin^-^, IL-7R^-^, Sca-1^-^, c-Kit^+^), GMP (granulocyte-monocyte progenitor, Lin^-^, IL-7R^-^, Sca-1^-^, c-Kit^+^, CD16/32^+^, CD34^+^), CMP (common myeloid progenitor, Lin^-^, IL-7R-, Sca-1^-^, c-Kit^+^, CD16/32^-^, CD34^+^), MEP (megakaryocyte-erythrocyte progenitor, Lin^-^, IL-7R^-^, Sca-1^-^, c-Kit^+^, CD16/32^-^, CD34^-^), CLP (common lymphoid progenitor, Lin^-^, IL7-R^+^, c-Kit^mid^, Sca-1^mid^); and *in vivo*-differentiated populations: erythroblast (CD71/CD44^+^), granulocyte (Gr-1^+^), monocyte (CD11b^+^), eosinophil/neutrophil (Gr-1/CD11b^+^), T cell (CD4 or CD8^+^) and B cell (B220^+^) are depicted as fold change compared to BM-injected mice. n = 6-12 per group, \**P* <0.05; \*\**P* <0.01; \*\*\**P* < 0.001 by Student *t*-test.

To determine cell numbers required for hematopoietic repopulation in mice, we gradually lowered the cell number used for injection. 2.5×10^6^ HPC^LSKs^ sufficed to allow for an 80% survival of the animals for a period of at least 8 months, after which the experiment was terminated. Injection of 1×10^6^ HPC^LSKs^ did not induce long-term survival but significantly prolonged the lifespan of lethally irradiated animals (median survival: 51 days compared to 8.5 days) (Supplementary Fig. S2b and S2c). These experiments led us to conclude that HPC^LSKs^ possess the ability for long-term replenishment of the hematopoietic system.

### Generation of leukemic HPC^LSKs^ as a model for leukemic stem cells (LSCs)

LSCs differ from the bulk of leukemic cells and possess the ability for self-renewal. To establish LSC models, we infected HPC^LSKs^ with a retrovirus encoding for oncogenes either inducing myeloid (BCR/ABL^p210^, MLL–AF9, Flt3-ITD;NRas^G12D^) or lymphoid (BCR/ABL^p185^) leukemia (Fig. 3a). Analysis of signaling pathways in the GFP^+^ leukemic lines showed that the cells faithfully reflected the signaling patterns downstream of the respective oncogene. As described, BCR/ABL predominantly induced phosphorylation of CRKL and STAT5^61,64^. Flt3-ITD;NRas^G12D^ was associated with a pronounced JAK2, STAT5, AKT and ERK signaling activation^65^ and MLL-AF9 upregulated c-MYC^66^ (Fig. 3b). In the presence of SCF and IL-6, transformed HPC^LSKs^ retained the expression of stem cell markers. A small fraction of the cells differentiated and upregulated the respective lineage markers. BCR/ABL positive LSCs were able to grow cytokine independently, whereas other oncogenes are shown with SCF (Fig. 3c-d). Except for MLL-AF9, all oncogenes tested formed growth factor-independent colonies in methylcellulose gel (Supplementary Fig. S3a).

**Figure 3:**
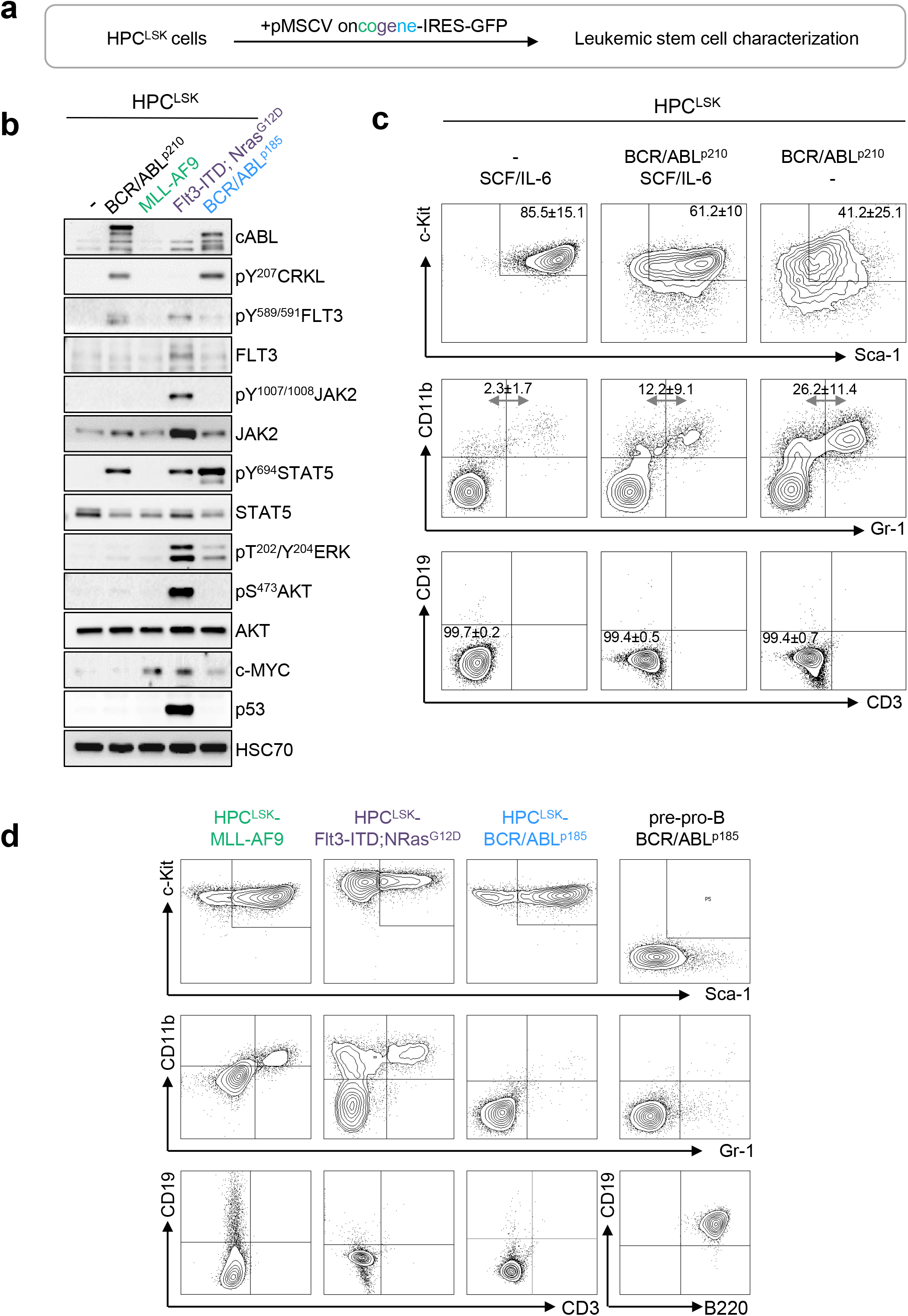
Successful generation of leukemic HPC^LSK^ cell lines with various oncogenes. (**a**) Experimental design: HPC^LSK^ cell lines were retrovirally transduced with different oncogenes. (**b**) Immunoblot showing increase of CRKL, FLT3, JAK2, STAT5, ERK, and AKT phosphorylation and upregulation of cABL, c-MYC and p53 in transformed HPC^LSK^ cells compared to untransformed (-) cells to the corresponding oncogenes. HSC70 serves as a loading control. Representative blot from at least three independent experiments is shown. (**c**) Flow cytometry analysis of untransformed and BCR/ABL^p210^ transformed HPC^LSK^ cells in IMDM/SCF/IL-6 and SCF/IL-6-deprived medium (IMDM). After one month in culture, HPC^LSK^ BCR/ABL^p210^ cells show reduced expression of stem cell markers (c-Kit, Sca-1) and differentiate into myeloid(CD11b, Gr-1), but not lymphoid (CD19, CD3) cells as indicated by the numbers in quadrants. The data are expressed as mean±SD of 3 independent measurements. (**d**) Representative flow cytometry plots of LSK (upper panel), myeloid (middle panel) and lymphoid staining (lower panel) of MLL-AF9 (in the presence of SCF and IL-6), Flt3-ITD;Nras^G12D^ and BCR/ABL^p185^ transformed HPC^LSK^ and pre-pro-B BCR/ABL^p185^ cell lines in the absence of SCF and IL-6.

To determine their leukemic potential *in vivo*, transformed HPC^LSKs^ were injected intravenously (*i.v*.) into NSG mice (Fig. 4a, left). HPC^LSKs^ BCR/ABL^p185^ inflicted disease within 12 days, followed by HPC^LSKs^ BCR/ABL^p210^ and HPC^LSKs^ Flt3-ITD;NRas^G12D^ which succumbed to disease within 50 days. The longest disease latency was observed upon injection of HPC^LSKs^ MLL-AF9 which induced disease after three months (Fig. 4a, right). All diseased animals displayed elevated WBC counts, blast-like cells in the blood and suffered from splenomegaly (Fig. 4b, 4d, Supplementary Fig. S4a). GFP^+^ transformed HPC^LSK^ cells were detected in the blood, spleen and BM of the diseased mice (Fig. 4c). HPC^LSKs^ BCR/ABL^p210^, HPC^LSKs^ MLL-AF9 and HPC^LSKs^ FLT3/NRas^G12D^-injected animals suffered from myeloid leukemia with an average of 92% CD11b^+^ cells, whereas HPC^LSKs^ BCR/ABL^p185^-injected NSGs developed predominantly GFP^+^ B cells with a percentage mean of 32% of CD19^+^ cells (Fig. 4e, Supplementary Fig. S4b-d). These experiments determine HPC^LSK^ cells as a valid model system studying leukemogenesis *in vivo* downstream of several oncogenic drivers.

**Figure 4:**
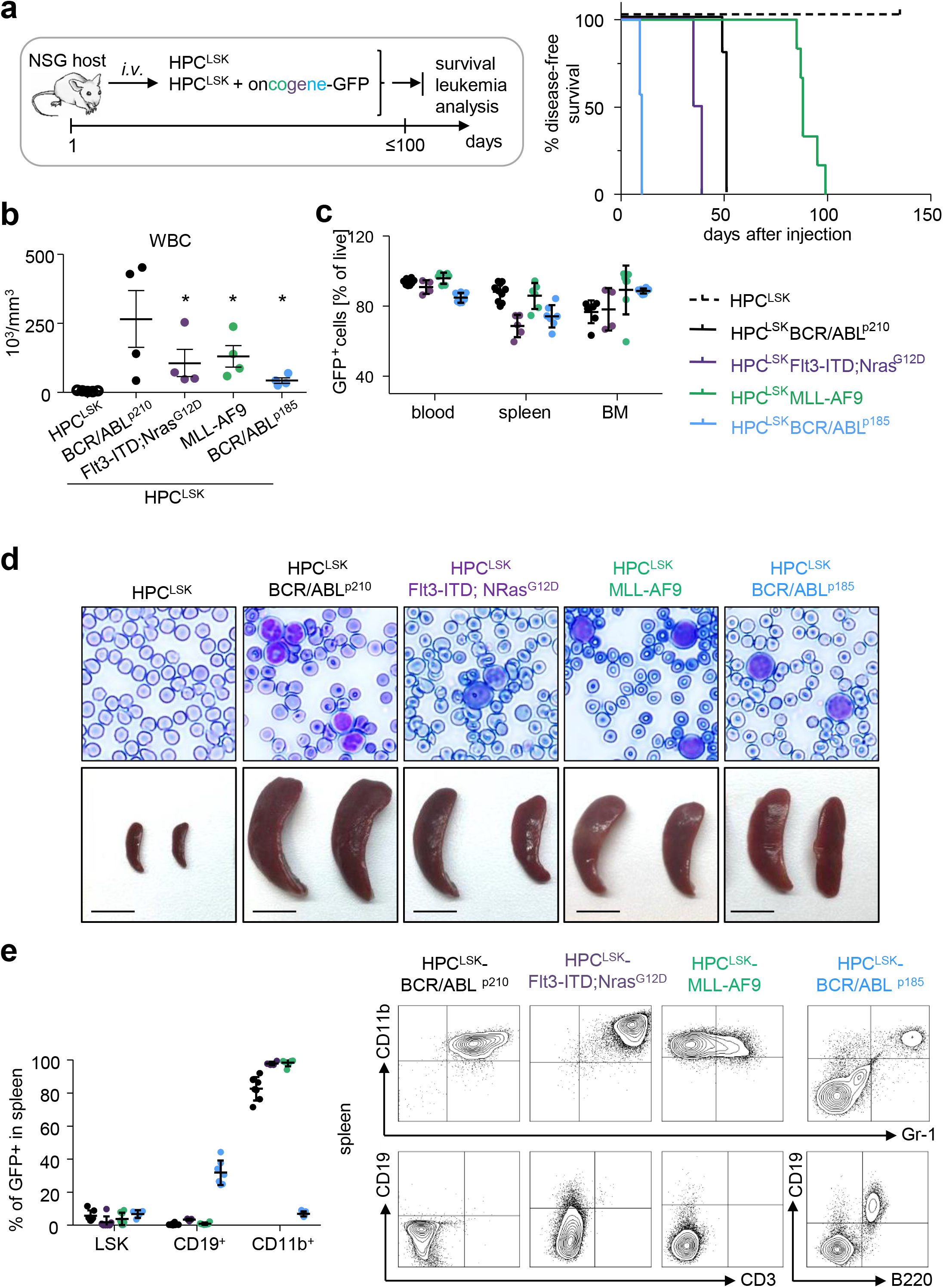
*In vivo* lymphoid and myeloid leukemia model. (**a**) Left: Schematic representation of the experimental setup; Oncogene-expressing HPC^LSK^ cell lines were injected *i.v*. in NSG recipients and moribund mice were analyzed. Healthy HPC^LSK^-injected animals were sacrificed and examined after 150 days. Right: Disease-free survival following *i.v*. injection of 2×10^6^ HPC^LSK^ BCR/ABL^p210^ (n=9), or 5×10^6^ HPC^LSK^ MLL-AF9 (n=7), HPC^LSK^ Flt3-ITD;NRas^G12D^ (n=5) and HPC^LSK^ BCR/ABL^p185^ (n=9) cells compared to injection of 5×10^6^ non-transformed HPC^LSK^ cells (n=5). (**b**) WBC count of moribund mice, One-way ANOVA (Kruskal-Wallis test) with Dunn’s Multiple Comparison Test, \**P*<0.05. Data are presented as mean±SEM. (**c**) Detection of transformed GFP^+^ HPC^LSK^ cells (with the respective oncogene) in blood, spleen and BM of diseased NSG recipients. Data represent mean±SD in 4-8 mice/group. (**d**) Top: Representative blood smears from transformed HPC^LSK^-injected mice shows leukocytosis with circulating blasts (hematoxylin-eosin, original magnification x400). Bottom: Macroscopic view of representative spleens from transformed HPC^LSK^-injected recipient mice compared to non-transformed HPC^LSK^-injected mice, n≥5. Scale bar: 1 cm. (**e**) Left: Quantification of the transformed GFP^+^ LSKs and differentiated cells (CD19^+^ B cells and CD11b^+^ myeloid cells) by flow cytometry in spleens of diseased NSG recipient mice. Error bars represent the mean±SD, n=4-8 per oncogene. Right: Representative flow cytometry plots for myeloid (CD11b and Gr-1) and lymphoid (CD19 and CD3 or B220) cells of spleens of the diseased mice injected with different oncogene-expressing HPC^LSKs^.

### HPC^LSKs^ from a transgenic mouse strain – *Cdk6^-/-^* HPC^LSKs^

CDK6 plays a key role as a transcriptional regulator for HSC activation and its function extends to LSCs^49^. To gain insights into distinct functions of CDK6 in HSCs/LSCs, we generated HPC^LSK^ cell lines from *Cdk6^-/-^* transgenic mice^60^. CDK4 does not compensate for the loss of CDK6 in those lines (Supplementary Fig. 5a). *Cdk6^-/-^* HPC^LSKs^ grow under normal HPC^LSK^ culture conditions albeit with a reduced cell proliferation and slightly increased apoptosis when compared to wild type HPC^LSKs^ (Fig. 5a, Supplementary Fig. 5b). 5×10^6^ *Cdk6^+/+^* or *Cdk6^-/-^* HPC^LSKs^ were equally well capable to rescue lethally irradiated mice for up to 60 days (data not shown).

**Figure 5:**
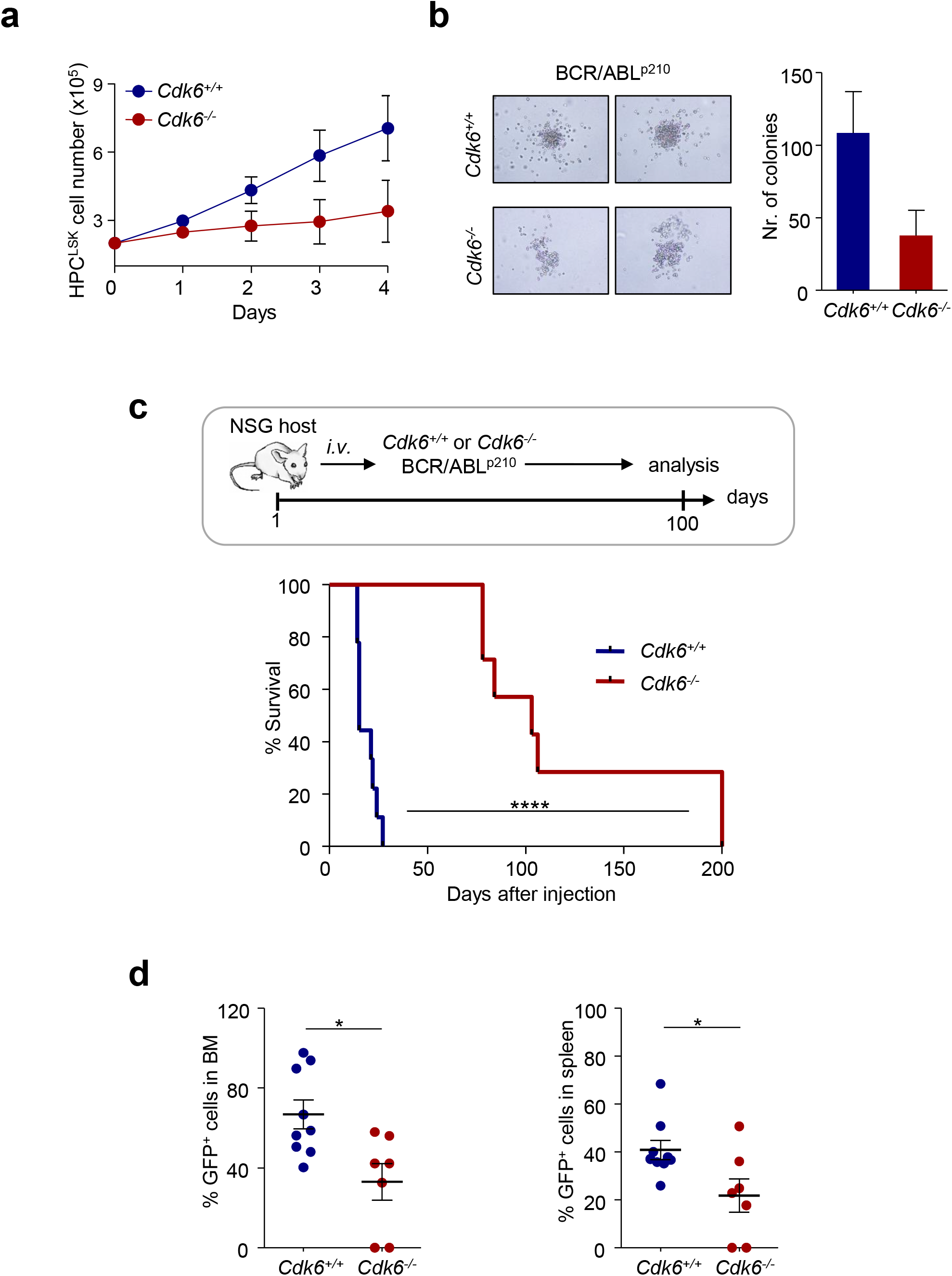
Generation of HPC^LSK^ lines from *Cdk6^-/-^* mice. (**a**) Cell proliferation curve of *Cdk6^+/+^* and *Cdk6^-/-^* HPC^LSK^ lines. Data are presented as mean±SEM of 3 different cell lines per genotype. (**b**) Colony formation assay of *Cdk6^+/+^* and *Cdk6^-/-^* BCR/ABL^p210^ HPC^LSKs^. Representative macroscopic images of colonies formed within 7 days in semi-solid methylcellulose gels without cytokines are depicted. Data are presented as mean±SEM of two independent experiments with 2-3 different cell lines per genotype. (**c**) Top: Schematic representation of the experimental setup; Bottom: *Cdk6^+/+^* and *Cdk6^-/-^* BCR/ABL^p210^ HPC^LSKs^ have been injected *i.v*. in NSG recipient mice. Disease-free survival following *i.v*. injection of 1×10^6^ *Cdk6^+/+^* (n=9, 3 different cell lines per genotype) and *Cdk6^-/-^* BCR/ABL^p210^ HPC^LSKs^ (n=7, 3 different cell lines per genotype). Statistical differences were calculated using the log-rank test (****, P < 0.0001); (**d**) Quantification of BCR/ABL^p210^ GFP^+^ cells by flow cytometry in BM and spleen of diseased NSG recipient mice. Error bars represent mean±SEM (n=7-9 per group, 3 different cell lines; \**P* <0.05 by Student *t*-test).

In a murine CML model BCR/ABL^p210^ *Cdk6^-/-^* BM cells induced disease significantly slower and with a drastically reduced disease phenotype^49^. To investigate whether this phenotype can be recapitulated with HPC^LSKs^, we generated *Cdk6^+/+^* and *Cdk6^-/-^* BCR/ABL^p210^ HPC^LSKs^ by retroviral infection. Irrespective of the presence of CDK6, BCR/ABL^p210^ HPC^LSKs^ grow in the absence of any cytokine and retain the expression of LSK markers (Supplementary Fig. 5c). In line with murine CML models, *Cdk6^-/-^* BCR/ABL^p210^ HPC^LSKs^ form fewer growth-factor independent colonies when compared to *Cdk6^+/+^* controls 7 days after plating, yet the difference did not reach significance (Fig. 5b)^49^. BCR/ABL^p210^ HPC^LSK^-derived colonies displayed Gr-1 and CD11b marker expression. However, *Cdk6^-/-^* BCR-ABL^p210^ HPC^LSKs^ show a trend to higher Gr-1 and lower CD11b expression compared to wild type (Supplementary Fig. S5d). To study the leukemic potential of *Cdk6^-/-^* BCR/ABL^p210^ HPC^LSKs^ *in vivo*, we injected 1*10^6^ cells *i.v*. into NSG mice. *Cdk6^+/+^* BCR/ABL^p210^ HPC^LSKs^ inflict disease within 14 days with severe signs of leukemia, including splenomegaly (Fig. 5c, Supplementary Fig. S5e). In contrast, *Cdk6^-/-^* BCR/ABL^p210^ HPC^LSKs^ failed to induce disease within this time period and only two thirds of the mice started to show signs of disease around 80 days after injection whereas one third of the animals did not develop any sign of leukemia within 7 months. Analysis of diseased mice show a reduced infiltration of *Cdk6^-/-^* BCR/ABL^p210^ HPC^LSKs^ into the BM and spleen, the percentage of BCR/ABL^p210^ GFP^+^ cells is comparable to *Cdk6^+/+^* control cells (Fig. 5d, Supplementary Fig. S5f). These results underline the crucial role of CDK6 in BCR/ABL^p210^ LSCs and verify the potential of our novel cellular HPC^LSK^ system to charter leukemic phenotypes.

### CDK6 dependent transcript alterations

To study CDK6-dependent gene regulation in untransformed and BCR/ABL^p210^ transformed HPC^LSKs^, we performed RNA-Seq analysis. Untransformed HPC^LSKs^ lacking CDK6 show an altered gene regulation with 1335 genes up- and 661 genes down-regulated when compared to *Cdk6^+/+^* HPC^LSKs^ (Fig. 6a). These differences decreased upon transformation; cytokine-independent BCR/ABL^p210^ HPC^LSKs^ showed 85 up- and 468 genes down-regulated in the absence of CDK6 compared to controls. Overall, 80% and 40% of genes found to be up- or downregulated in *Cdk6^-/-^* BCR/ABL^p210^ HPC^LSK^ cells were also de-regulated in *Cdk6^-/-^* untransformed HPC^LSK^ cells defining a transformation-independent gene signature downstream of CDK6 (Fig. 6B). Gene Ontology enrichment analyses of CDK6 dependent genes revealed an association with immune response, cell adhesion, cell death and myeloid cell differentiation irrespective of the transformation status (Fig. 6C). The differential gene expression in our murine BCR/ABL^p210^ HPC^LSK^ cells was compared to CDK6 associated gene expression changes in human CML samples. To do so, we stratified a dataset from 76 human CML patients into CDK6^high^ and CDK6^low^ samples based on quartile expression of CDK6 and subsequently calculated the differential gene expression. We identified 101 genes that are regulated in a CDK6–dependent manner in murine and human BCR/ABL^p210^ cells (Fig. 6D). In human and mouse CDK6 dependent deregulated genes belong to pathways pointing at apoptosis/stress response, cell differentiation and homing.

**Figure 6:**
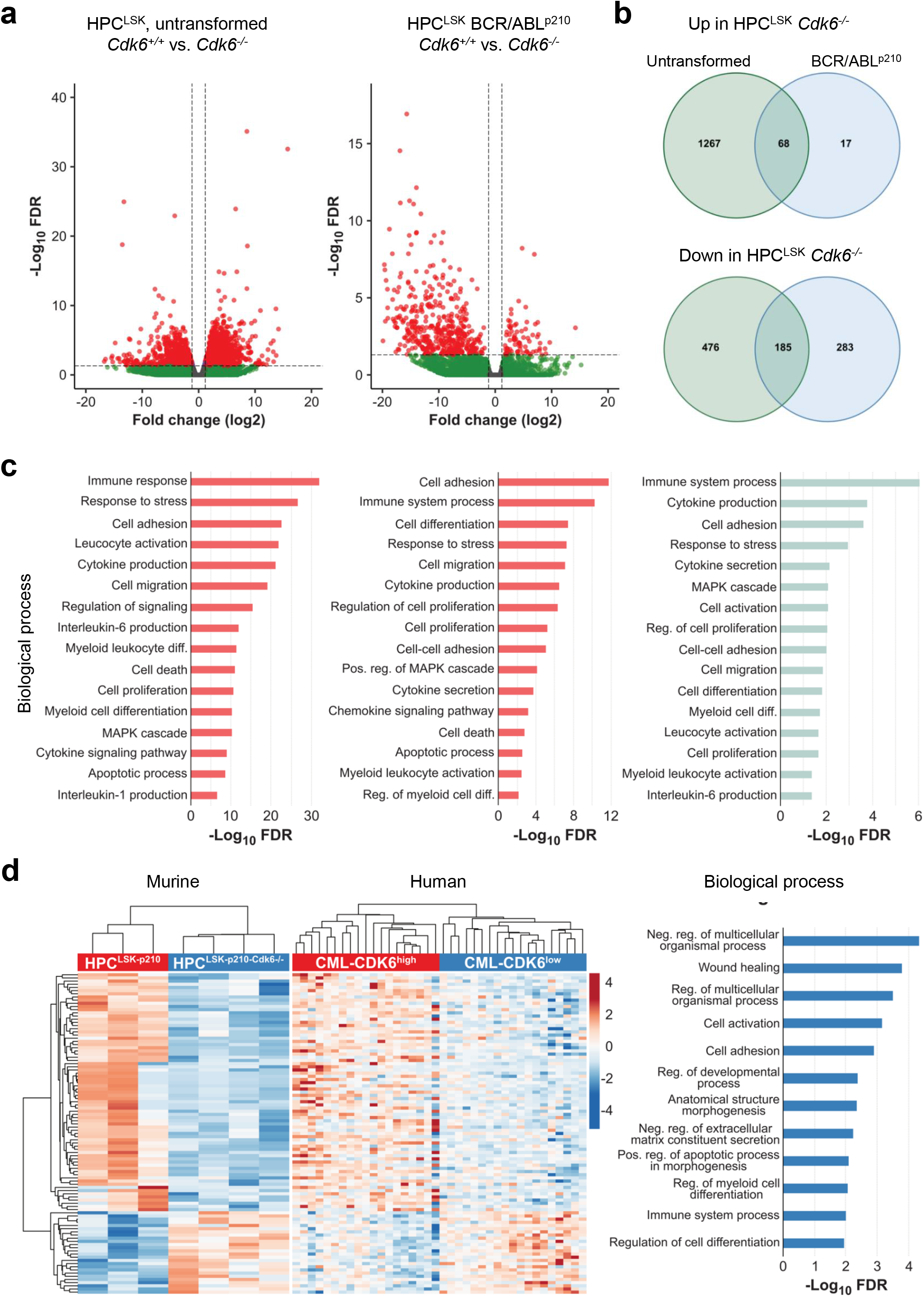
CDK6 dependent transcriptomic alterations. **(a)** Volcano plots summarizing Cdk6-mediated differential gene expression between untransformed (left) and BCR/ABL^p210^ (right) HPC^LSKs^. Each dot represents a unique gene, red dots indicate statistically significant deregulated genes (FDR<0.05 and FC±1.5). FDR, false discovery rate; FC, fold change. **(b)** Venn diagrams showing overlaps between upregulated genes (upper panel) or downregulated genes (lower panel) in untransformed and *Cdk6^-/-^* BCR/ABL^p210^ HPC^LSKs^. **(c)** Gene Ontology enrichment analyses of Cdk6 regulated genes in untransformed (left) and BCR/ABL^p210^ (middle) HPC^LSKs^ and of commonly Cdk6 regulated genes in these cell types (right). **(d)** Heatmaps summarizing expression of 101 genes which are commonly regulated in a CDK6-dependent manner in murine and human BCR/ABL^p210^ cells. Each row represents a unique gene and each column represents a unique sample. Colors range from blue (low expression) to red (high expression). Results from Gene Ontology enrichment analyses of these genes are shown in the bar chart (right).

### Validation of CDK6 dependent pathways in LSCs

In line with the deregulated pathways in human and mouse resulting from the RNA-Seq analysis, we recently demonstrated that CDK6 regulates apoptosis during BCR/ABL^p185^ transformation^57^. To validate this aspect in our HPC^LSK^ system, we serum starved *Cdk6^+/+^* and *Cdk6^-/-^* BCR/ABL^p210^ HPC^LSKs^ for 90 minutes and performed an apoptosis staining by flow cytometry (Fig. 7a). As expected, *Cdk6^-/-^* BCR/ABL^p210^ HPC^LSKs^ showed increased response to stress.

**Figure 7:**
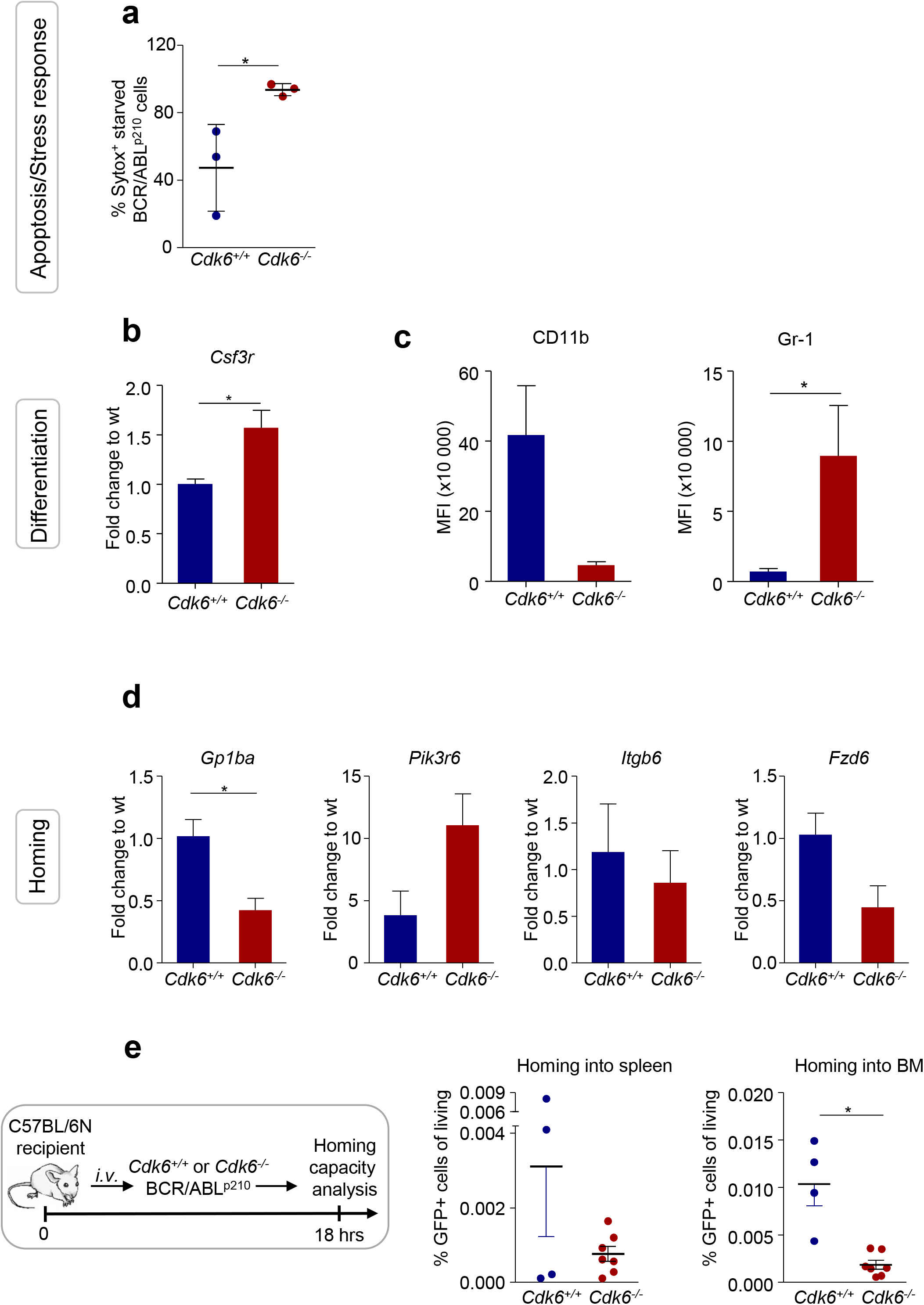
CDK6 is required for homing to the bone marrow of BCR/ABL^p210^ HPC^LSKs^. **(a)** Sytox staining for apoptotic cells of BCR/ABL^p210^ HPC^LSKs^ starved for 90 min in 0,5% FCS medium. Numbers represent mean±SD (n=3 different cell lines per genotype; \**P* <0.05 by Student *t*-test.). **(b)** qPCR validation of RNA-Seq data of the target gene *Csf3r* (mean± SEM; n=3 different cell lines per genotype; \**P* <0.05 by Student *t*-test.) **(c)** Mean fluorescence intensity (MFI) of myeloid markers (CD11b, Gr-1) of BCR/ABL^p210^ HPC^LSKs^ (mean± SEM; n=3 different cell lines per genotype; \**P* <0.05 by Student *t*-test) **(d)** Validation of 4 selected genes (*Gp1ba, Pik3r6, Itgb6, Fzd6*) found deregulated in GO analysis of the RNA-Seq experiment by qPCR and nested qPCR. (mean± SEM; n=3 different cell lines per genotype; \**P* <0.05 by Student *t*-test) **(e)** Upper part: Experimental scheme of BCR/ABL^p210^ HPC^LSKs^ homing assay in wild type recipient mice; Bottom: Percentage of BCR/ABL^p210^ GFP^+^ HPC^LSKs^ in spleen and BM detected by flow cytometry are shown. (mean± SEM; n=4-7 per group, two-three independent cell lines, \**P* <0.05 by Student *t*-test)

In addition to apoptosis, cell differentiation was one of the most significant deregulated pathways detected by the transcriptome analysis. Colonies from *Cdk6^-/-^* BCR/ABL^p210^ HPC^LSKs^ showed a bias to the granulocytic direction by increased Gr-1 expression (Supplementary Fig. S5c). In the RNA-Seq analysis and validated by qPCR, *Csf3r*, an essential receptor for granulocytic differentiation, is upregulated in *Cdk6 ^-/-^* BCR/ABL^p210^ cells compared to controls (Fig. 7b). Further, cytokine independent *Cdk6^-/-^* BCR/ABL^p210^ HPC^LSKs^ show increased mean fluorescence intensity (MFI) levels of Gr1 and reduced MFI levels of CD11b compared to *Cdk6^+/+^* controls (Fig. 7C). Together, these data demonstrate that loss of CDK6 shows an advantage for granulocytic differentiation.

Last but not least, the reduced percentages of *Cdk6^-/-^* BCR/ABL^p210^ HPC^LSKs^ in the BM and spleen upon *i.v*. injection (Fig. 5d) together with the RNA-Seq analysis point towards a hampered homing capacity of *Cdk6^-/-^* BCR/ABL^p210^ HPC^LSK^ cells. We validated several deregulated genes found in the transcriptome analysis which can be linked to homing by qPCR analysis (Fig. 7D) and performed an *in vivo* homing assay. To do so, we injected 1*10^6^ BCR/ABL^p210^ HPC^LSKs^ with and without CDK6 into aged and gender matched female C57BL/6N mice and profiled the number of BCR/ABL^p210^ GFP^+^ cells after 18h in the BM and spleen by flow cytometry. *Cdk6^-/-^* BCR/ABL^p210^ HPC^LSKs^ showed a significantly diminished homing capability to the BM compared to *Cdk6^+/+^* BCR/ABL^p210^ HPC^LSKs^.

Taken together, the validated data describes essential roles of CDK6 in LSCs and supports the strong reliability of our murine cellular system. Moreover, we here describe a prominent function for CDK6 in regulating BCR/ABL^p210^ leukemic cell homing.

## DISCUSSION

Functional and molecular studies on hematopoietic and leukemic stem cells have provided numerous insights into the mechanisms of hematopoietic diseases. However, progress is restricted by the limited availability of hematopoietic stem/progenitor cells and the difficulty of *in vitro* culturing. We present a robust procedure to generate an unlimited source of functional mouse HSC/HPC lines called HPC^LSK^ that possess characteristics of MPPs and can serve as a source of lymphoid and myeloid LSC lines. HPC^LSKs^ are multipotent cells that retain lymphoid and myeloid differentiation potential and can repopulate lethally irradiated mice without supporter BM cells. More than 90% of HPC^LSKs^ are Lin^-^/c-Kit^+^/Sca-1^+^ and express CD34, CD48 and CD150, which is characteristic of MPP2. They also express CD41, which marks cells at the embryonic AGM that constitute the myelo-erythroid and myelo-lymphoid branchpoint in early hematopoiesis^67–68^. The transcriptome of the cells most closely resembles that of MPP1 cells, which correspond to the earliest proliferating stem/progenitor cell.

Our approach is robust and simple and requires no co-culture system or feeder layer and no extensive amounts of cytokines. We have established more than 50 distinct HPC^LSK^ cell lines with an efficiency of 100%, using either mouse strains of various genetic backgrounds or transgenic mice as a source. HPC^LSK^ cells can be genetically modified by retroviral transduction or CrispR/Cas9 technologies, so are a versatile tool in HSC and LSC research. Our method is based on the enforced expression of *Lhx2*, a transcription factor for mouse HPC immortalization^39,41–42,46–47^. Improvements to the original protocol include FACS sorting of LSKs to avoid 5-FU treatment, the use of serum low-media with a defined cocktail of cytokines, pre-coating of plates to avoid adherence–induced myeloid differentiation and the maintenance of high HPC^LSK^ cell density^4,41,47,69–75^. *Lhx2*-immortalized HPCs have been reported to induce a transplantable myeloproliferative disorder resembling human chronic myeloid leukemia in long-term engrafted mice^76^. We did not observe this even after long-term repopulation in lethally irradiated Ly5.1 or in immunosuppressed NSG mice. The difference probably stems from our use of sorted LSKs instead of total BM to overexpress *Lhx2*, as the myeloid disorder may originate from a more differentiated myeloid progenitor. We have used HPC^LSKs^ as a source to generate leukemic stem cells and obtained leukemic HPC^LSK^ lines harboring BCR/ABL, MLL-AF9 and Flt3-ITD;NRas^G12D^ oncogenes. Removal of SCF and IL-6 *in vitro* induced myeloid differentiation, indicating that the self-renewal program depends on the presence of low-level cytokines and downstream signaling events that are provided *in vivo* by the BM niche.

The cell cycle kinase CDK6 is a transcriptional regulator and is particularly important in hematopoietic malignancies. In HSCs, its actions are largely independent of its kinase activity. It is essential for HSC activation in the most dormant stem cell population under stress situations, including transplantation and oncogenic stress. The impact of CDK6 extends to leukemic stem cells, as BCR/ABL^p210^-transformed BM cells fail to induce disease *in vivo* in the absence of CDK6. To investigate how CDK6 drives leukemogenesis in progenitor cells, we generated *Cdk6^-/-^* HPC^LSKs^ from *Cdk6*-deficient mice and transformed them with BCR/ABL^p210^. The absence of CDK6 was associated with a reduced incidence of leukemia and with significantly delayed disease development, thereby mimicking the effects seen in primary bone marrow transplantation assays. RNA-Seq and subsequent pathway analysis show deregulated stress response, cell adhesion and apoptotic processes/cell death in the absence of CDK6. This result is consistent with our recent observations that CDK6 antagonizes p53 responses and regulates survival. In the absence of CDK6, hematopoietic cells need to overcome oncogenic-induced stress by mutating p53 or activating alternative survival pathways, as in the case of Cdk6-deficient JAK2^V617F^ positive LSKs. Another featured shared by CDK6-deficient JAK2^V617F+^ LSKs and CDK6-deficient BCR/ABL HPC^LSK^ is an altered cytokine secretion, as revealed by pathway enrichment analysis in both systems. HSCs show homing and cell adhesion, which allow them to migrate to the bone marrow and replenish hematopoietic lineages^77^. GO pathway analysis revealed deregulated cell adhesion and cell migration pathways in HPC^LSK^ cell lines and in human patient samples. Our bioinformatic data show that loss of CDK6 from transformed cells leads to a significantly reduced capacity to home to the bone marrow, which slows the onset of leukemic disease. The common CDK6 dependent gene signature between BCR/ABL^p210^ HPC^LSKs^ and human CML patient samples underlines the translational relevance of our model system. A large subset of CDK6 regulated genes is also found in patients, which we could validate with specific assays using our BCR/ABL^p210^ HPC^LSKs^. The data strengthen our confidence in our murine cellular system and show that results from HPC^LSK^ experiments can be translated to the human situation. HPC^LSK^ lines thus represent a quick and simple alternative to the lymphoid progenitor Ba/F3 or the myeloblast-like 32D cells to explore the potential transforming ability of mutations found in hematopoietic malignancies.

## Author contributions

ED and IMM designed and conducted experiments, collected and analyzed data. TB, BM and IM collected and analyzed data. RG, MZ and GH performed bio-informatical analysis. LC was involved in conception and design of the study, contributed essential material and reviewed the manuscript. KK designed and supervised experiments. AHK reviewed the manuscript and supervised experiments. VS designed and supervised the study, VS, ED, IMM, BM and KK wrote the manuscript.

## Acknowledgements

We thank P. Kudweis, S. Fajmann, M. Ensfelder-Koparek and P. Jodl for excellent technical support and M. Dolezal for critical discussion of bioinformatical analysis. We thank the Biomedical Sequencing Facility (BSF) at CeMM for NGS library preparation, sequencing and related bioinformatics analyses.

## Funding

This work was supported by the European Research Council (ERC) under the European Union’s Horizon 2020 research and innovation programme grant agreement No 694354. This work was supported by the Austrian Science Foundation (FWF) via grants to K.K. (P 31773).

## Conflict of interest

The authors declare that they have no conflicts of interest.

## SUPPLEMENTARY FIGURE LEGENDS

**Supplementary Figure 1:**
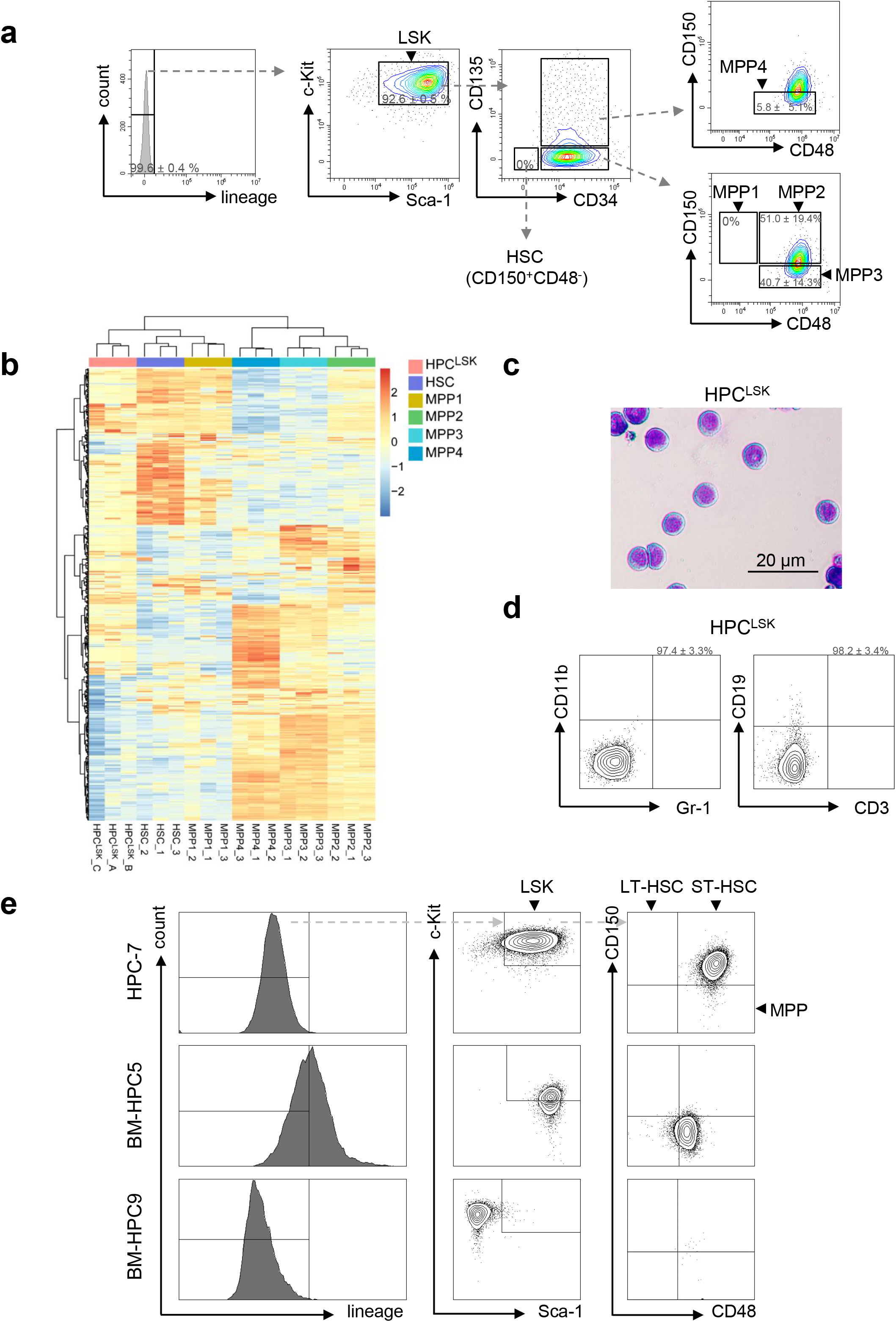
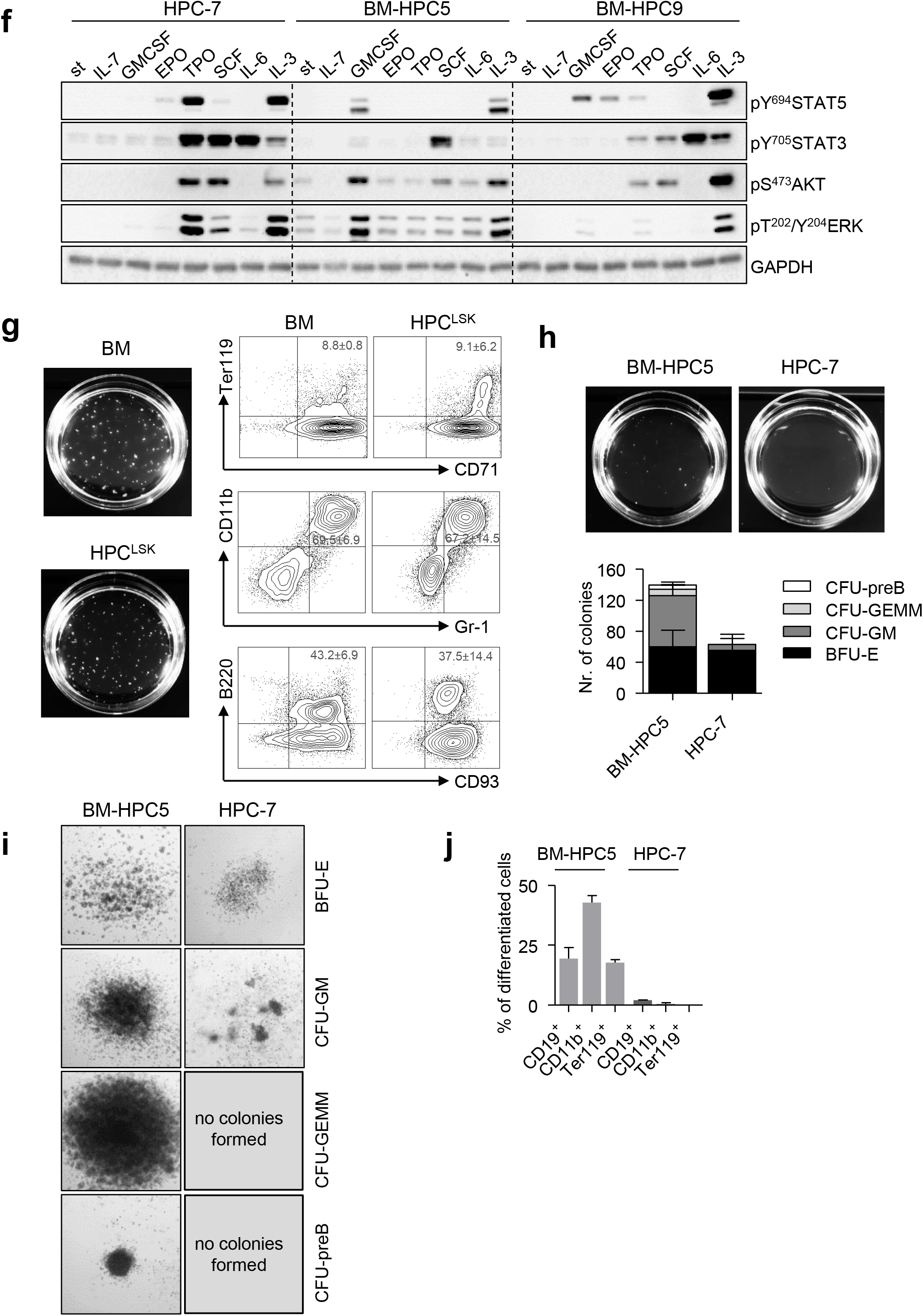
Comparison of HPC^LSK^ cell line with other *Lhx2*-immortalized murine hematopoietic progenitor cell lines. (**a**) Representative plots of HPC^LSK^ cell line for Sca-1, c-Kit, CD135, CD34, CD150 and CD48. LSK population that contains hematopoietic stem cells: HSC, MPP1, MPP2, MPP3 and MPP4; One representative example is depicted. All data represent mean±SD of five HPC^LSK^ cell line establishments. (**b**) Heatmap of batch-corrected top500 variance genes of RNA-expression profiles of HPC^LSKs^ (n=3) compared to murine HSCs. (**c**) Light microscopy picture of cultured HPC^LSKs^ (Cytospin stained with haematoxylin/eosin), scale bar depicts 20 μm. (**d**) Representative myeloid (CD11b, Gr-1) and lymphoid (CD19, CD3) flow cytometry plots of HPC^LSK^ cell line. All data represent mean±SD of four HPC^LSK^ cell line establishments. (**e**) HSC staining of ES-derived HPC-7, BM-HPC5 and BM-HPC9 cell lines. Cells were gated on lineage-negative, c-Kit and Sca-1 positive, and stained for CD150 and CD48. (**f**) Immunoblot analysis of lysates from HPC-7, BM-HPC5 and BM-HPC9 cell lines. Samples were starved for 3 h, then treated with IL-7, GMCSF, EPO, TPO, SCF, IL-6 or IL-3 (100 ng/ml each) for 15 min and blotted for phosphorylated STAT5, STAT3, AKT and ERK. GAPDH serves as a loading control. Representative blot from three independent experiments. st, starved. (**g**) Representative macroscopic pictures and flow cytometry analysis of colonies formed by BM and HPC^LSK^ cells in cytokine cocktail supplemented methylcellulose gel. The numbers in quadrants indicate the percentage of erythroid Ter119/CD71^+^, myeloid CD11b/Gr-1^+^ and lymphoid B220/CD93^+^ B cells after 10 days. The data are expressed as the mean±SD of 3 independent experiments. (**h**) Macroscopic, number of colonies and (**i**) microscopic images of colonies on methylcellulose gels formed by BM-HPC5 and HPC-7 cells 10 days after a cytokine cocktail treatment (EPO, GMCSF, IL-7, SCF, IL-6, IL-3). Primitive erythroid progenitor (BFU-E), multipotential progenitor (CFU-GEMM), lymphoid (CFU-preB) and myeloid (CFU-GM) progenitor colonies were counted. Seeding density of 1 250 cells/35-mm-dish. Error bars represent mean+SD, n=3. (**j**) Flow cytometry analysis of HPC-7 and BM-HPC5 cells upon colony formation in cytokine cocktail-supplemented methylcellulose gel. The bars indicate the percentage CD19^+^ B cells, CD11b^+^ myeloid cells and Ter119^+^ erythroid cells after 10 days. Data represent mean±SD, n=3.

**Supplementary Figure 2:**
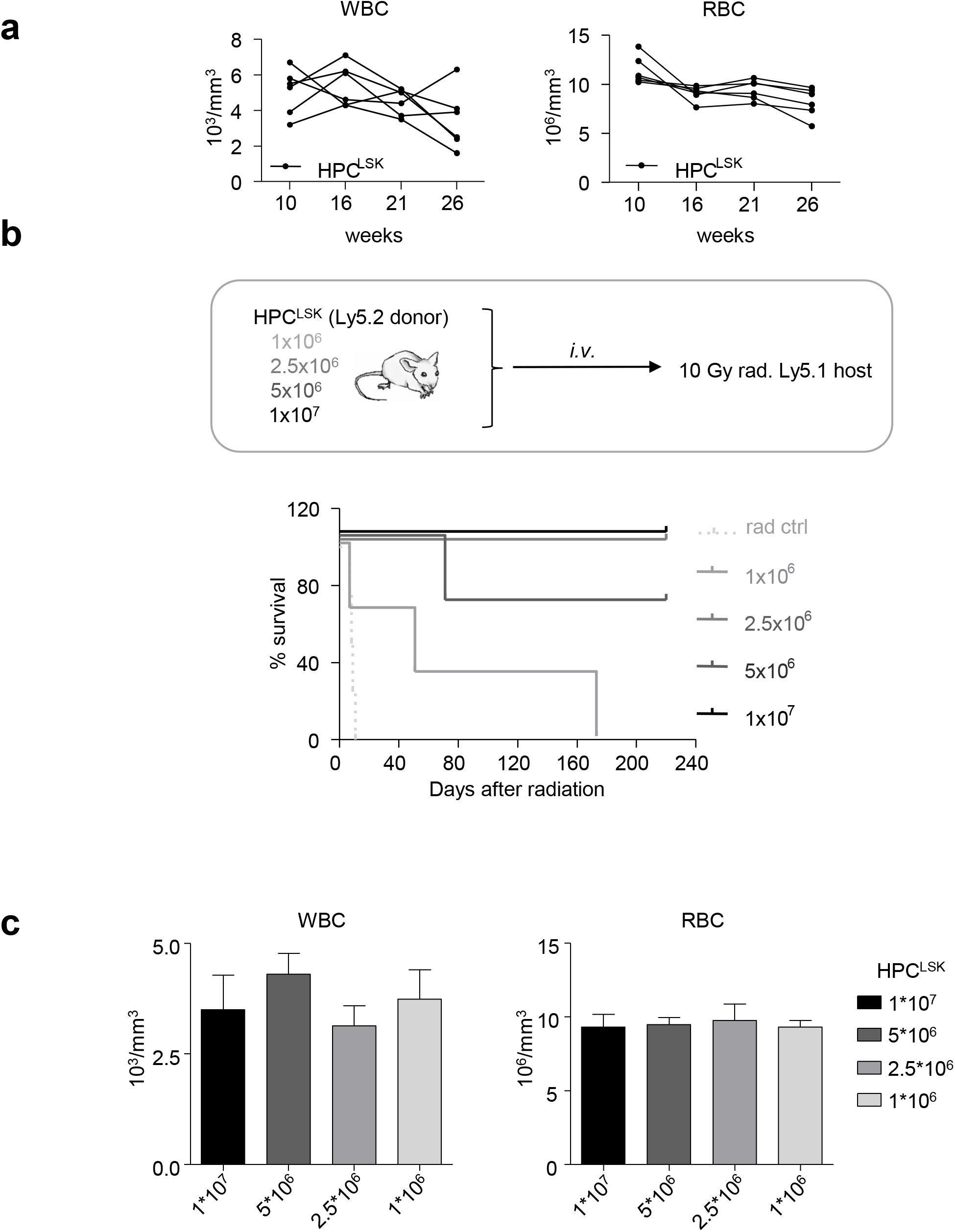
*In vivo* differentiation of HPC^LSK^ cell lines. (**a**) WBC and RBC values of 5 mice were followed after lethal irradiation (10 Gy) and 1×10^7^ HPC^LSKs^ injection for 26 weeks. (**b**) Top: Experimental scheme: Ly5.1^+^ recipient mice were lethally irradiated 24 h prior to *i.v*. injection of different (1-10×10^6^) numbers of Ly5.2^+^ HPC^LSK^ cells. Bottom: Long-term survival, n = 4/group; (**c**) Absolute numbers of WBC and RBC of mice injected with 1-10×10^6^ HPC^LSK^ cells were compared 40 days after transplantation. Bars represent mean±SD, n≥3.

**Supplementary Figure 3:**
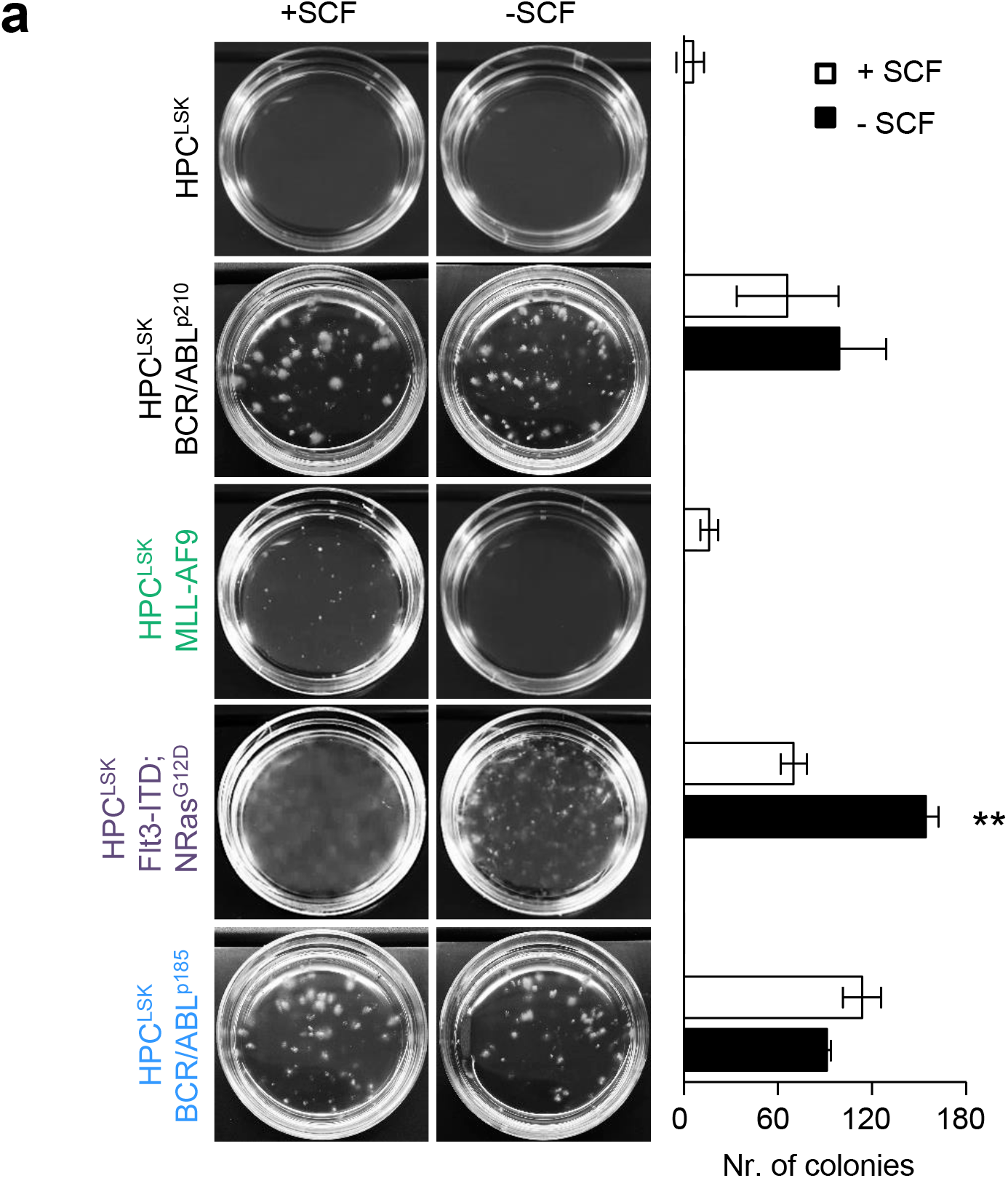
Oncogenic transformation of HPC^LSK^ cell lines. (**a**) Representative macroscopic pictures of dishes and colony counts of methylcellulose gels formed by oncogene-transformed HPC^LSK^ cells with or without SCF 10 days after seeding, density of 1 250 cells/35-mm-dish. Data represent mean±SD, n=3, Two-way ANOVA with Bonferroni post-test, **P<0.01.

**Supplementary Figure 4:**
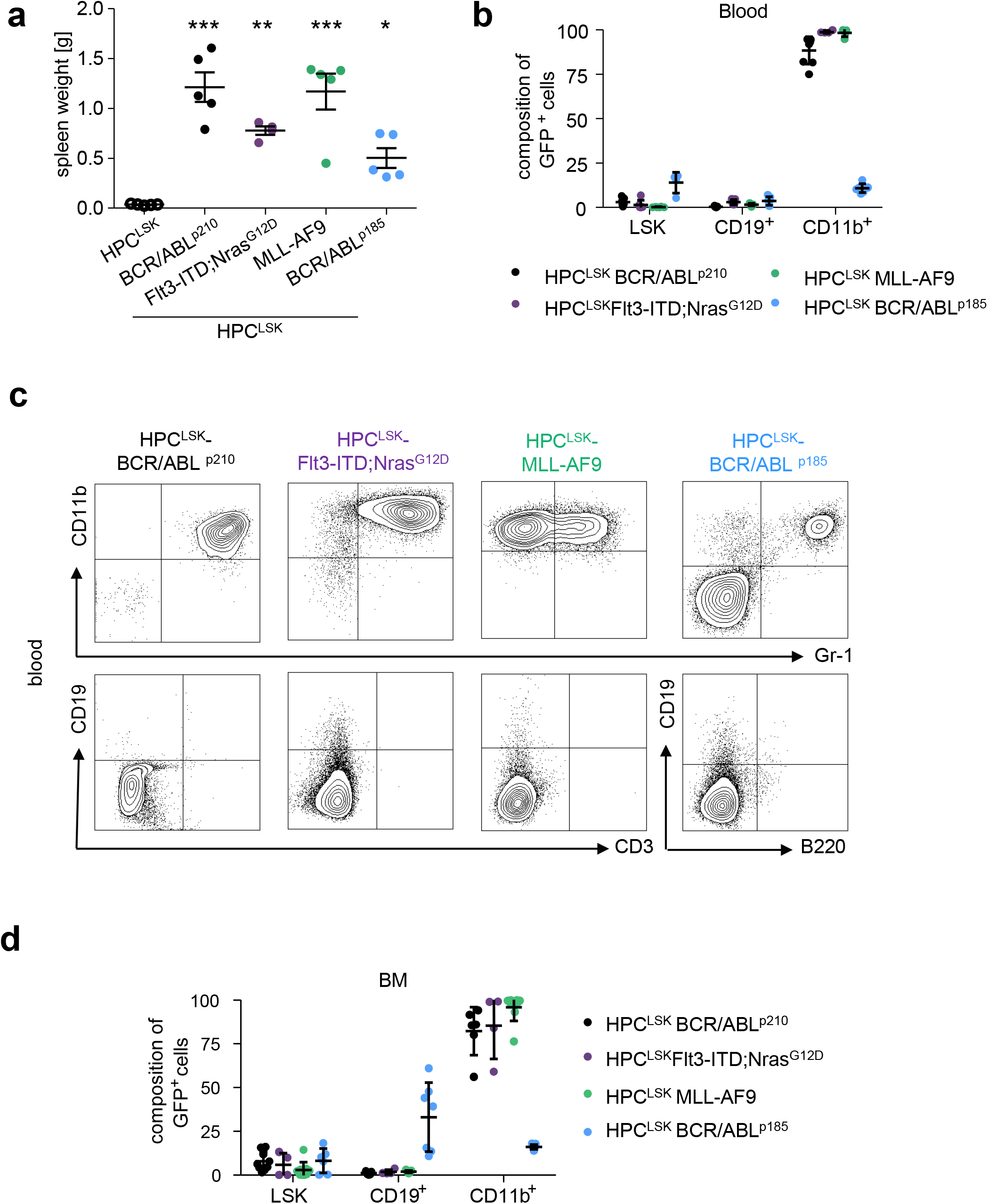
Characterization of transformed HPC^LSK^-induced leukemia. (**a**) Spleen weights of moribund transformed HPC^LSK^ recipients. One-way ANOVA with Bonferroni’s Multiple Comparison Test (*P<0.05, **P<0.01, ***P<0.001). Error bars represent mean±SEM. (**b**) Quantification of transformed GFP^+^ LSKs and differentiated cells (CD19^+^ B cells and CD11b^+^ myeloid cells) by flow cytometry in the blood of diseased NSG recipient mice. Error bars represent the mean±SD. n=4-8 per oncogene. (**c**) Representative blood flow cytometry plots (myeloid CD11b and Gr-1 and lymphoid CD19 and CD3 or B220 staining) of the diseased mice injected with different oncogene-expressing HPC^LSKs^. (**d**) Quantification of transformed GFP^+^ LSKs and differentiated cells (CD19^+^ B cells and CD11b^+^ myeloid cells) by flow cytometry in the BM of diseased NSG recipient mice. Error bars represent mean±SD. n=4-8 per oncogene.

**Supplementary Figure S5:**
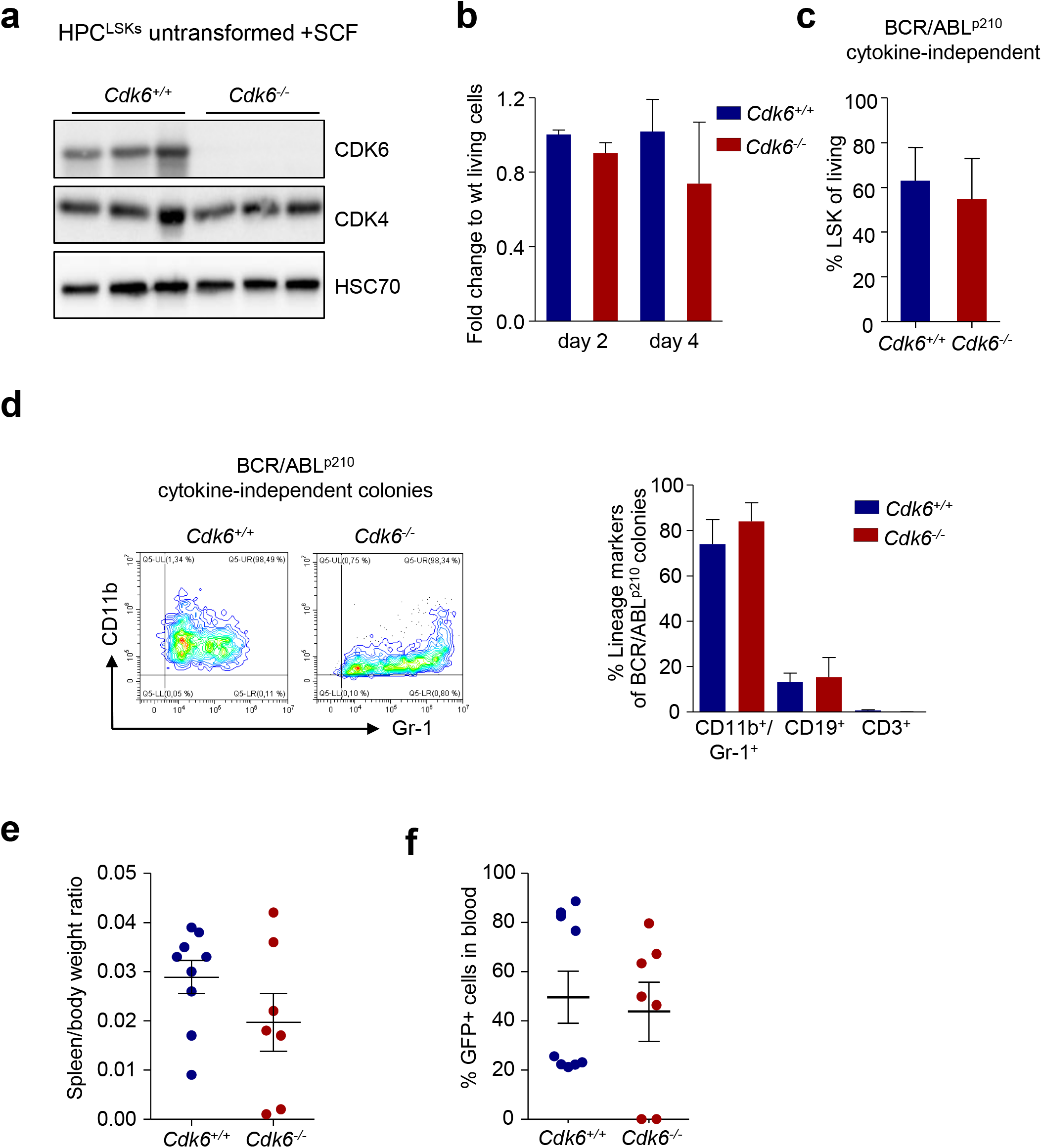
Validation of transgenic HPC^LSK^ *Cdk6^-/-^* cell line under physiological conditions and transformation. (**a**) Immunoblot for CDK6 and CDK4 of three untransformed *Cdk6^+/+^* and *Cdk6^-/-^* HPC^LSK^ lines. HSC70 serves as a loading control. (**b**) Fold change of % living cells of the cell proliferation curve of *Cdk6^+/+^* and *Cdk6^-/-^* HPC^LSKs^ at day 2 and 4. The data is presented as mean±SEM of 3 independent cell lines/genotype. (**c**) Analysis of transformed GFP^+^ LSKs (Gr1/CD11b^+^ myeloid cells) by flow cytometry after cytokine removal for 2 weeks. The data are presented as mean±SEM of 2-3 independent cell lines/genotype and two independent experiments. (**d**) Left: Representative flow cytometry plots of BCR/ABL^p210^ HPC^LSK^ colonies in the absence of SCF and IL-6 for myeloid lineage markers (CD11b and Gr-1). Right: Flow cytometry analysis of *Cdk6^+/+^* and *Cdk6^-/-^* BCR/ABL^p210^ HPC^LSK^ colonies after 7 days in semi-solid methylcellulose culture for the myeloid lineage (CD11b, Gr-1), B-cell lineage (CD19) and T-cell lineage (CD3). Data represent mean±SEM, n=4-8/genotype. (**e**) Spleen and body weights were measured on the day of sacrifice and spleen/body weight ratio was calculated. The data is presented as mean±SEM, n=7-9/genotype. (**f**) Quantification of transformed GFP^+^ LSKs by flow cytometry in blood of diseased NSG recipient mice. Error bars represent mean±SEM. n=7-9 per group; *p*= <0.055 by Student *t*-test.

## Notes

### Competing Interest Statement

The authors have declared no competing interest.

